# Single cell transcriptome profiling of the Atlantic cod immune system

**DOI:** 10.1101/2020.01.31.926410

**Authors:** Naomi Guslund, Monica Hongrø Solbakken, Kjetill S. Jakobsen, Shuo-Wang Qiao

**Affiliations:** University of Oslo

## Abstract

The Atlantic cod’s unusual immune system, entirely lacking the Major Histocompatibility class II pathway, has prompted intriguing questions about what mechanisms are used to combat bacterial infections and how immunological memory is generated. Here, we examine the diversity of 8,180 spleen cells and peripheral blood leukocytes by single cell RNA sequencing. Unbiased transcriptional clustering revealed eleven distinct immune cell signatures. Resolution at the single cell level enabled characterisation of the major cell subsets including the cytotoxic T cells, B cells, erythrocytes, thrombocytes, neutrophils and macrophages. Further, we describe for the first time rare cell subsets which may represent dendritic cells, natural killer-like cells and a population of cytotoxic cells expressing GATA-3. We propose putative gene markers for each cluster and describe the relative proportions of each cell type in the spleen and peripheral blood leukocytes. By single cell analysis, this study provides the most detailed molecular and cellular characterization of the immune system of the Atlantic cod so far.

## Introduction

Teleosts, accounting for nearly half of all extant vertebrates (Ravi and Venkatesh 2018), demonstrate an extraordinary level of diversity within their habitat, morphology, physiology, behaviour and in the genetic repertoire of their immune system (Zhu, Nie et al. 2013, Solbakken, Torresen et al. 2016, Solbakken, Voje et al. 2017). Whole genome sequencing of the Atlantic cod (*Gadus morhua*) and other Gadiform species revealed that genes encoding Major Histocompatibility class II (MHCII) were missing, along with the absence of the entire CD4+ T cell component of the adaptive immunity (Star, Nederbragt et al. 2011, Malmstrom, Matschiner et al. 2016). For the first time, it was demonstrated that this classical immune pathway can no longer be considered the hallmark of vertebrate immunity (Malmstrom, Matschiner et al. 2016, Parham 2016, Solbakken, Voje et al. 2017). The evolutionary events leading to the loss of MHCII are not known, but several putative past biological scenarios have been suggested (Star and Jentoft 2012, Solbakken, Voje et al. 2017). Further peculiarities mark the Atlantic cod immune system, including extreme expansion of MHC class I (MHCI) genes as well as gene losses and expansions within the innate immune system (Star, Nederbragt et al. 2011, Solbakken, Rise et al. 2016, Solbakken, Torresen et al. 2016, Torresen, Brieuc et al. 2018), a low to modest response by specific antibodies following pathogen exposure but a consistently high level of natural IgM (Solem and Stenvik 2006, Magnadottir, Gudmundsdottir et al. 2009). Since the Atlantic cod and codfishes demonstrate such an interesting immune system, understanding the cells and genes involved in immune responses in this species is an evolutionary intriguing question with respect to the workings of an immune system that naturally lacks CD4+ T cells, and in providing insights into the flexibility of the vertebrate immune system. Lastly, a better grasp of the cod immune system would be beneficial for improved management of cod stock and potential cod aquaculture where infectious disease is a challenge.

Traditionally, immune cells are characterized using the unique combination of cell markers present on the cell surface, but a lack of specific antibody-based reagents makes this approach difficult. Advances in next-generation sequencing technologies allow for a closer examination of biological systems without the need for existing antibodies. The use of single cell RNA sequencing (scRNA-seq) makes it possible to examine the global mRNA content of thousands of individual cells, facilitating a more detailed characterisation. Cell types can be clustered computationally using bioinformatic tools according to their transcriptional activity, and by analysing the transcriptional fingerprint in comparison to an annotated genome cell types and functions can be assigned.

A combination of microscopy and *in vitro* functional studies have already identified some immune cell types in Atlantic cod, while whole genome and transcriptome sequencing have led to the identification of putative cell markers. The functional assignment and cell markers of cytotoxic CD8+ T cells (CD8, TRGC1, TNFSF11, EOMES, TCR, CD3), B cells (IGLC2, IgM, CD79), natural killer (NK)-like cells (LITR/NITR, B3GAT1), cells with granules and perforin activities (PERF1, UNC13D), monocytes/macrophages (IL34) and neutrophils (MPO) have been described to some extent within the Atlantic cod (Fischer, Utke et al. 2006, Øverland, Pettersen et al. 2010, Star, Nederbragt et al. 2011, Solbakken, Jentoft et al. 2019a, Solbakken, Jentoft et al. 2019b). Cells found in the blood and organs of related teleost species might also be expected in the Atlantic cod, including thrombocytes (Nagasawa, Nakayasu et al. 2014), nonspecific cytotoxic cells (NCCs) (Evans, Leary et al. 1998, Shen, Stuge et al. 2002, Shen, Stuge et al. 2004) and dendritic cells (DCs) (Lugo-Villarino, Balla et al. 2010, Bassity and Clark 2012). However, the relative proportion of these verified and putative immune cell subsets is unknown, and there is no overall assessment of the cellular functions. Further, cell markers to date are largely based on knowledge about the mammalian immune system, and further cell type characterization by means of single cell RNA sequencing could lead to the development of antibodies towards new and specific Atlantic cod cell markers.

In this study, we used single-cell RNA-seq to investigate the gene expression of over 8,000 individual peripheral blood leukocytes (PBLs) and spleen cells from Atlantic cod. Combined with conventional morphological microscope studies, we identified 13 cell subsets, of which 11 are likely immune cell populations, and their putative markers. Thus, we have provided for the first time a systematic overview of the relative frequencies of these cell populations in blood and spleen. This study provides the most detailed molecular and cellular characterization of the immune system of the Atlantic cod so far.

## Methods and Materials

### Atlantic cod sampling

The Atlantic cod from one single breeding family (bred from 1 female and 2 males) were originally supplied as juveniles by NOFIMA (Tromsø, Norway) where there is a national breeding program for cod. The cod were reared at the NIVA Research Facility at Solbergstrand (near Oslo), Norway. The rearing and sampling are performed according to animal welfare regulations and approved by the Norwegian authorities (FOTS ID 12336). The water temperature is kept at an average of 8°C, with salinity at 34 PSU, fish are fed with Skretting cod pellets and checked twice a day and exposed to 12/12 hr illumination. Blood and spleen samples were taken from two specimens of non-vaccinated, 2-year-old Atlantic cod; 1 male (fish 1, 47cm, 0.99kg) and 1 female (fish 2, 52cm, 1.77kg) were killed within seconds of capture by cranial concussion. Neither fish showed visible signs of infection on skin, gills, fins or internally. Blood samples were collected from the *vena caudalis* with heparinized syringes. The spleens were removed and placed in Leibovitz L-15þ (L-15 (BioWhittaker) (adjusted to 370 mOsm by adding 5% (v/v) of a solution consisting of 0.41 M NaCl, 0.33 M NaHCO3 and 0.66% (w/v) D-glucose)) and transported on ice. Spleen cell suspensions were obtained by gently forcing the tissue through a cell strainer (Falcon, 100 µm). Blood samples of 0.7ml were diluted in L-15þ to a total volume of 5 ml. The blood cell suspensions were placed on discontinuous Percoll gradients (3 ml 1.070 g/ml overlaid with 2.5 ml 1.050 g/ml) and centrifuged for 40 min at 400xg and 4°C. A peripheral blood leukocyte (PBL) fraction was collected from the interface of the two Percoll densities, including the downward density layer, and washed twice by diluting the suspension in L-15þ and centrifuging at 300xg for 7 min at 4°C. All cells were kept in regular microcentrifuge tubes to minimise any cell loss and kept on ice at all times.

### Sorting of cell populations by flow cytometry

Spleen, blood and PBL suspensions were further separated into sub-populations on a FACS Aria II flow cytometer (Flow Cytometry Core facility at Oslo University Hospital) gated on the forward scatter (FSC, cell size) vs side scatter (SSC, granularity) plot. The sorted populations were examined by microscopy after cytospin and staining (supplementary Figure 1). Sub-populations of potential interest from the blood (B1, containing a large lymphocyte population), the spleen (S3, containing a large myeloid cell population) and from PBLs (P1, P2 and P3) as well as unsorted spleen and PBL cell suspensions were subjected to scRNA-seq using the Drop-seq protocol (Macosko, Basu et al. 2015). An overview of the samples can be seen in supplementary table 1.

**Figure 1.**
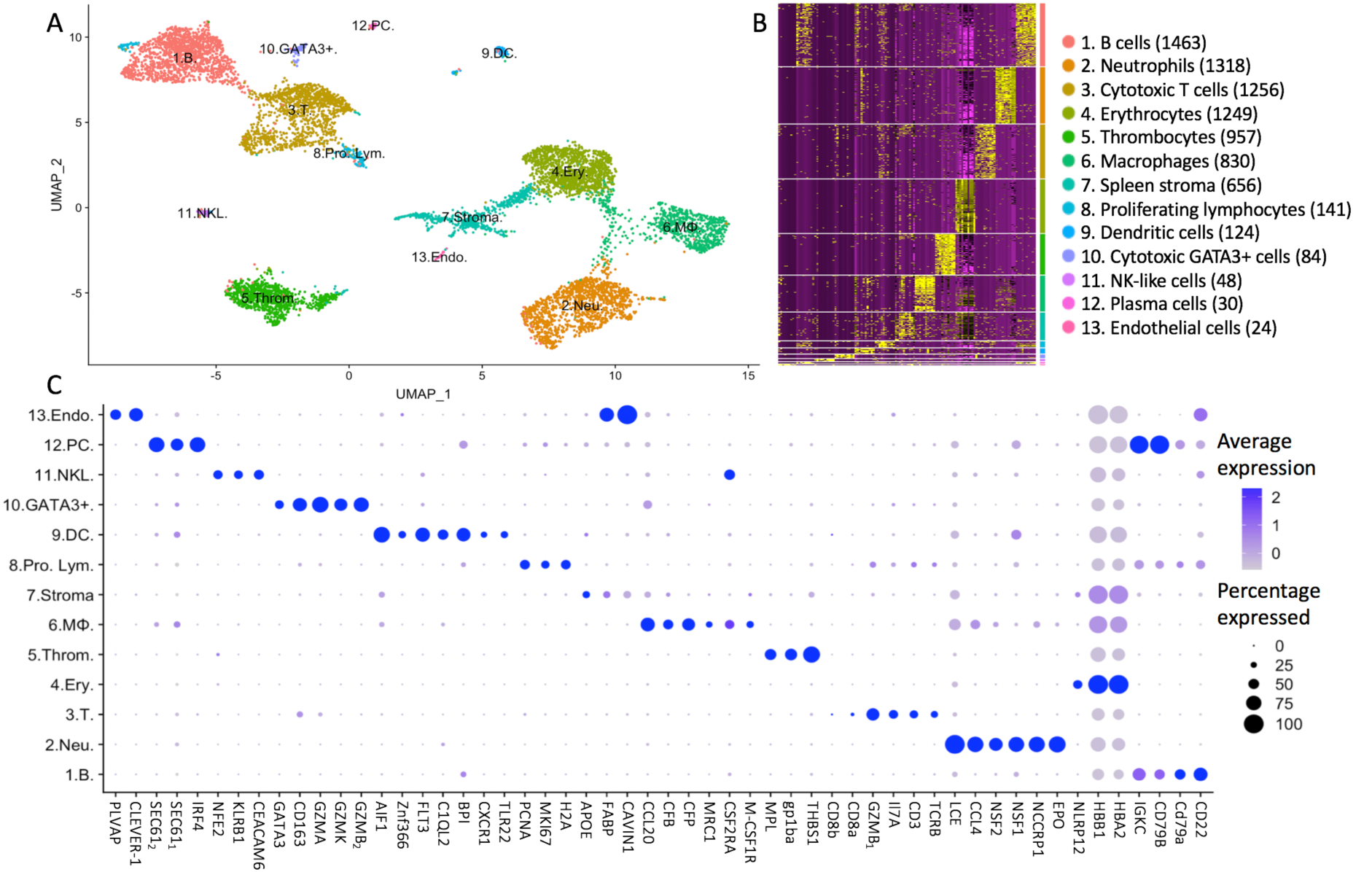
Unsupervised clustering of transcriptomic profiles of Atlantic cod immune cells demonstrates distinct cell populations. A: Visualization of cell types in two dimensions using UMAP. Putative cell cluster labels are based upon the differential gene expression of known markers in mammals, overall expression pattern and microscopy. B: Heatmap showing differential gene expression, with top 10 differentially expressed genes for each cluster shown (supplementary excel sheet). Each row represents a cell while each column represents a gene. The clusters are shown in descending population size, from B cells to endothelial cells respectively. The size of each cell population is shown in brackets. C: Dot plot showing the putative marker genes across the cell clusters. The size of the dot encodes the percentage of cells within a cluster expressing the gene, while the colour intensity encodes the average expression level of ‘expressing’ cells. *Two GZMB and SEC61 genes are identified (denoted _1_ and _2_ for clarity). Key: 1. B. are B cells, 2. Neu. are neutrophils, 3. T. are T cells, 4. Ery. are erythrocytes, 5. Throm. are thrombocytes, 6. MΦ. are macrophages, 7. Stroma. are spleen stromal cells, 8. Pro. Lym. Are proliferating lymphocytes, 9. DC. are dendritic cells, 10. GATA3+. are cytotoxic GATA3+ cells, 11. NKL. are natural-killer like cells, 12. PC. are plasma cells, 13. Endo. are endothelial cells.

### Staining of sorted cell populations

Immediately after sorting, 10,000 cells in 15 µl PBS buffer were added to a Cytospin carrier and subjected to centrifugation onto glass slides (80 g for 3 minutes). Slides were then air-dried and stained either next day or stored at −20C until fixation and staining. The slides were subjected to routine Hematoxylin-eosin (HE) staining. In short, slides of cells were briefly stained with hematoxylin and then by eosin solution, followed by alcohol dehydration. Some cells were also stained for peroxidase activity with the iVIEW DAB detection kit (Roche) according to the manufacturer’s instructions.

### scRNA-seq with Drop-seq protocol

The protocol and reagents used closely followed the protocol written by the McCarroll laboratory (Macosko and Goldman 2015), which is an amended version of the method used by Macosko *et al*., 2015. Briefly, the cell suspension was fed through a droplet generator (Dolomite, UK) that encapsulated a single cell and a barcoded bead in a water-in-oil droplet with a diameter of approximately 80 μm. All the beads contain a primer with a common “PCR handle” sequence to enable PCR amplification. Each individual bead contains 10^8^ primers with the same “cell barcode” but also contain unique molecular identifiers (UMIs), thus enabling the transcripts to be digitally counted at a later stage. A 30-bp oligo dT sequence for the capture of mRNAs is incorporated at the end of the primer. When a cell and bead are enclosed in a droplet the cell is lysed and the poly-dT sequences capture the released mRNA, forming single-cell transcriptomes attached to microparticles (STAMPs). The STAMPs are reverse-transcribed to make cDNA, amplified, and barcoded fragments generated by Tn5-mediated tagmentation. During the post-tagmentation PCR, unique sample barcodes were introduced in the adaptor primers so that samples from different cell populations could be multiplexed in the same sequencing library.

### Quantification of genes

The libraries were sequenced at the Norwegian Sequencing Centre (Oslo University Hospital), on the NextSeq500 platform with a 75 bp kit, high output mode, with paired end reads. 20 bp were sequenced in Read 1 using a custom sequencing primer (GCCTGTCCGCGGAAGCAGTGGTATCAACGCAGAGTAC) and 60 bp in Read 2 with the regular Illumina sequencing primer. We used the Drop-seq Core Computational Protocol (Namesh 2015) using STAR alignment to map the raw sequencing data to the most recent version of the Atlantic cod genome, gadMor3 (RefSeq accession GCF_902167405.1). A gene of interest, GATA-3, was present in gadMor2 (Torresen, Star et al. 2017) but missing in gadMor3, so the GATA-3 gene sequence was manually added to the gadMor3 assembly fasta file. Reads were then grouped by cell barcode and the unique molecular identifiers (UMIs) for each gene counted, resulting in a digital expression matrix showing the number of transcripts per gene per cell. From each sample, reads from the first 600-5000 STAMPs (depending on sample size) in decreasing number of reads were included into the next steps for filtering (Supplementary Figure 2). Further analysis was performed using R version 3.4.4.

**Figure 2.**
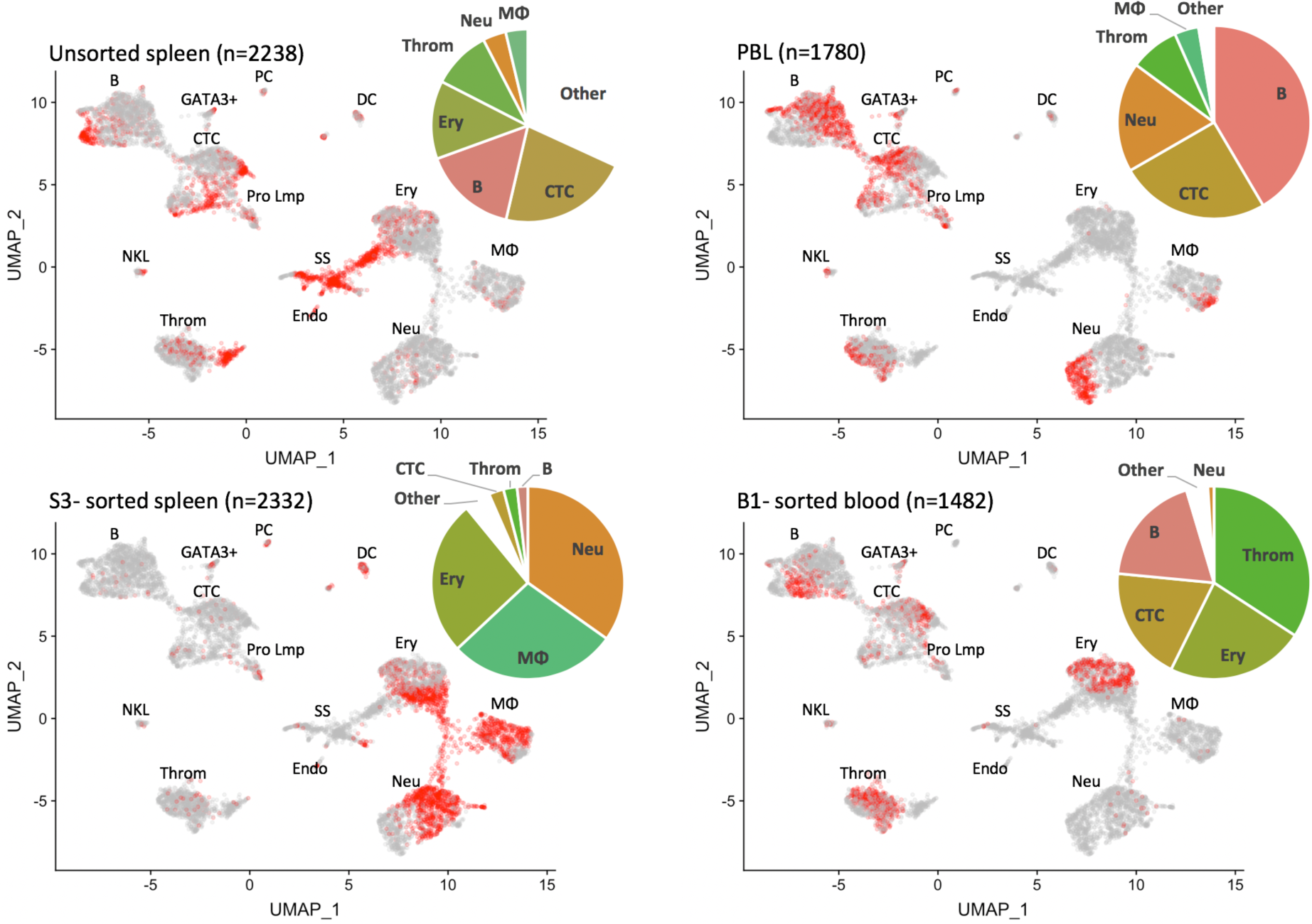
UMAP feature plots show the distribution of cells from Atlantic cod unsorted spleen, isolated peripheral blood mononuclear cells (PBLs) and sorted sub-populations. Population B1 (mostly lymphocytes and thrombocytes) is from the blood; S3 (mostly myeloid cells) is from the spleen. The number of cells per sample is given in brackets (n=). The cells from the named sample are shown in red. The pie charts show the percentage distribution of each cell type in the sample. Resting and proliferating T cells are grouped together, whereas resting and proliferating B cells as well as plasma cells are grouped as B cells. Key: 1. B. are B cells, 2. Neu. are neutrophils, 3. T. are T cells, 4. Ery. are erythrocytes, 5. Throm. are thrombocytes, 6. MΦ. are macrophages, 7. Stroma. are spleen stromal cells, 8. Pro. Lym. Are proliferating lymphocytes, 9. DC. are dendritic cells, 10. GATA3+. are cytotoxic GATA3+ cells, 11. NKL. are natural-killer like cells, 12. PC. are plasma cells, 13. Endo. are endothelial cells.

### Cell and gene selection

We followed the unsupervised clustering analysis tutorial (Butler, Hoffman et al. 2018) on the R package Seurat 3.0.2. The data matrix from all samples was merged to create one Seurat object. Cells with a gene count of fewer than 150 and a gene count of more than 1500, and cells with a total number of molecules of more than 4000 were filtered away in order to remove low-quality cells and possible cell multiplets (supplementary Figure 3). By excluding genes expressed in less than 5 cells (among the cells having passed the quality control), 15,273 genes across 8,180 cells was used in the study. An overview of sample origin, average mapping percentages, included cells, mapped transcripts and genes are shown in supplementary table 1.

**Figure 3.**
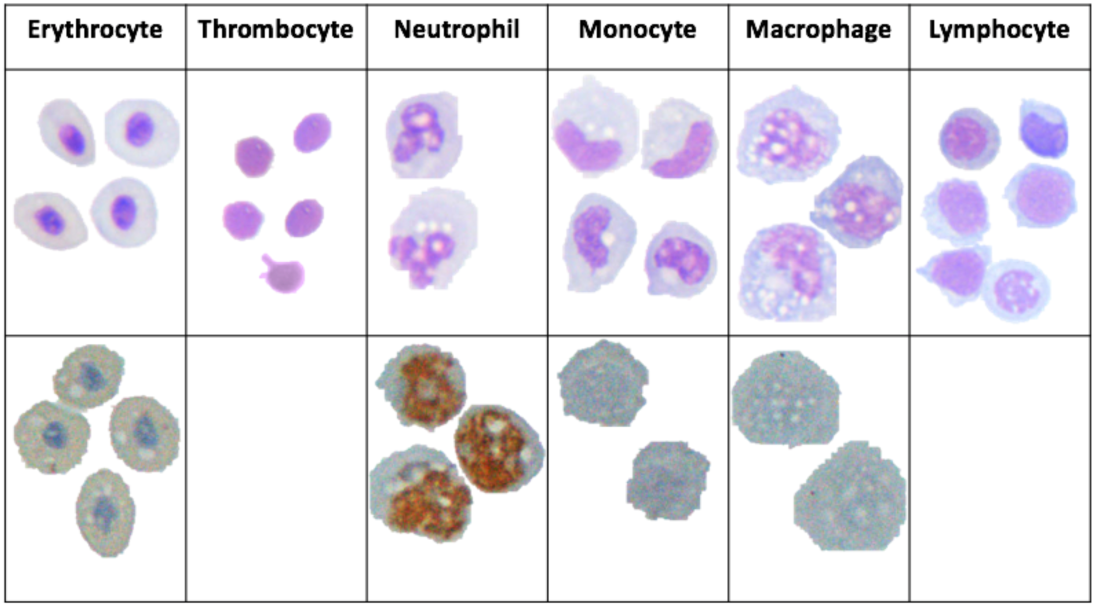
Microscopy images of Atlantic cod immune cells. Cells in the top panel are HE-stained (A) and the cells in the bottom panel are stained with peroxidase.

### Cell clustering and visualization

After “LogNormalize” and scaling (with a scale factor 10,000), we ran principal component analysis (PCA) on the expression of the top 2000 variable genes. We then used the FindCluster function in Seurat in order to cluster cells based on a shared nearest neighbor (SNN) modularity optimization result on the top 30 principal components (PCs) (supplementary figure 4). The resolution for clustering was 0.35. The cell clusters were visualized by the non-linear dimensional reduction method uniform manifold approximation and projection (UMAP).

**Figure 4.**
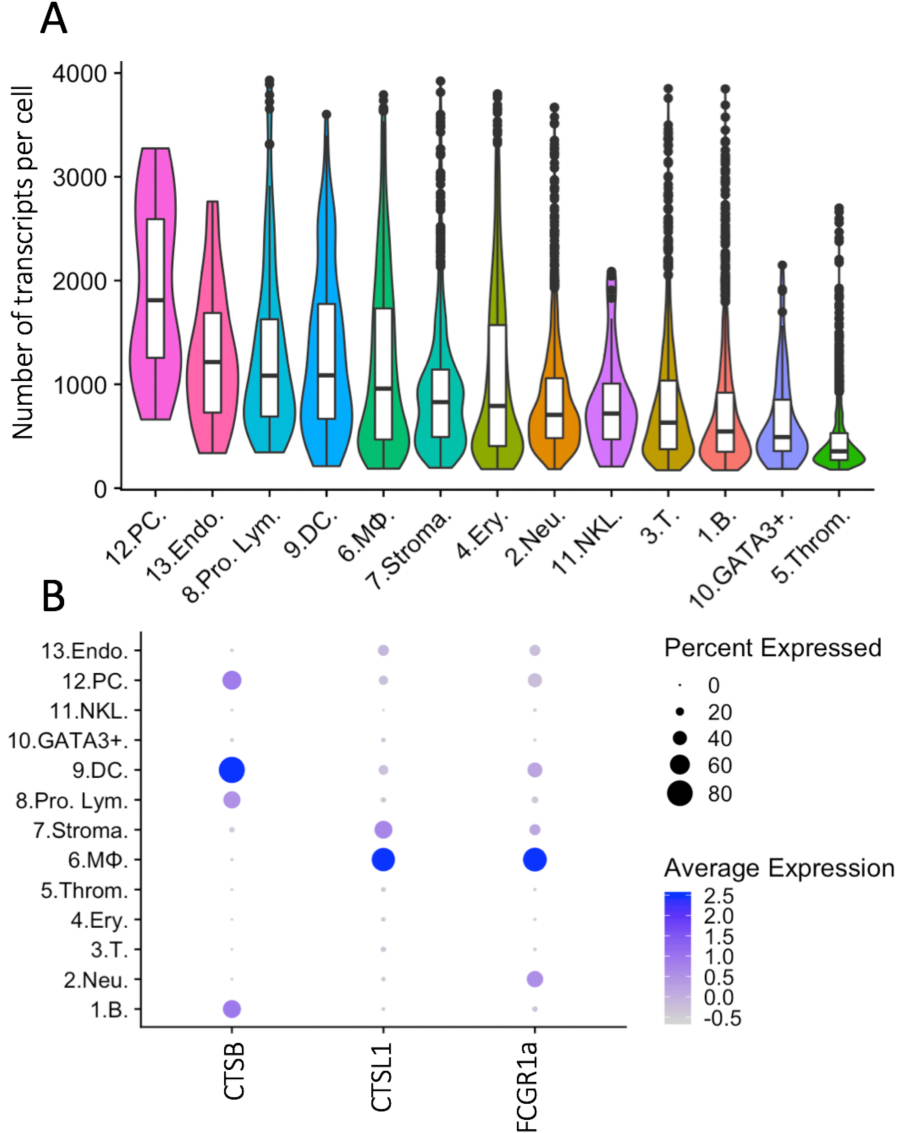
The overall transcriptional activity of each cell type and differential expression of selected genes. A) Violin plots showing the average number of transcripts per cell within the cluster. The overlaid boxplots show the mean and the 25% and 75% percentile of transcripts. B) Dot plot showing the expression of selected genes across the cell clusters. The size of the dot displays the percentage of cells within a class expressing the gene, while the colour intensity encodes the average expression level of ‘expressing’ cells. Key: 1. B. are B cells, 2. Neu. are neutrophils, 3. T. are T cells, 4. Ery. are erythrocytes, 5. Throm. are thrombocytes, 6. MΦ. are macrophages, 7. Stroma. are spleen stromal cells, 8. Pro. Lym. Are proliferating lymphocytes, 9. DC. are dendritic cells, 10. GATA3+. are cytotoxic GATA3+ cells, 11. NKL. are natural-killer like cells, 12. PC. are plasma cells, 13. Endo. are endothelial cells.

### Differentially expressed genes

All differential expression analyses in this study were performed using FindMarkers embedded in the Seurat package, which performs differential expression based on the non-parameteric Wilcoxon rank sum test. Adjusted p-values were calculated based on Bonferroni correction. In order to be counted as a differentially expressed gene, the gene must be expressed by a minimum of 25% of cells in the cluster. The most significantly differentially expressed genes for each cluster are listed in order of significance (supplementary excel sheet).

### Additional data from three wild-caught cod

In addition, similar analyses were carried out on unsorted blood and spleen samples from three other Atlantic cod in pilot studies. These samples were taken from wild-caught Atlantic cod from Oslo fjord, taken by direct sampling alongside the project “Monitoring cod health in the inner Oslo fjord”(Ulset 2018). Data from this pilot experiment accounts for an additional 15,758 genes across 3,744 cells (supplementary excel sheet and supplementary figure 6).

## Results

Following stringent quality control filtering, we derived a gene expression matrix of 15,273 genes across 8,180 cells (Supplementary Table 1). Visualization of cell types in two dimensions using Uniform Manifold Approximation and Projection (UMAP) (Figure 1A) reveals 13 cell clusters with distinct gene expression signatures, with cell cluster sizes ranging from 1,463 cells to the smallest cluster with only 24 cells. Overlapping distribution of cells from the two fish (Supplementary Figure 5) support the robustness of the clusters. Additionally, the clusters formed (supplementary figure 6) and the top differentially expressed genes expressed by each cluster (supplementary excel sheet), are similar to those produced in the pilot study.

To assign a cell identity to each cell cluster we performed differential gene expression analysis. Identified populations include a lymphocyte lineage, with B cells and plasma cells, T cells, cytotoxic GATA3^+^ cells and putative NK-like cells and a myeloid lineage that includes erythrocytes, thrombocytes, neutrophils and macrophages (Figure 1C). A putative DC population is also described. Additionally, some cells have been classified as spleen stroma and endothelial cells.

The largest cell cluster was identified as the B cell population based on CD22, CD79 and immunoglobulin genes such as IGKC. A small population of B cells, also expressing immunoglobulin genes and CD79, were identified as plasma cells or plasmablasts due to the expression of the transcription factor interferon regulatory factor 4 (IRF4) which controls plasma cell differentiation (Klein, Casola et al. 2006). These cells also show high expression of the transport protein SEC61, which mediates the transport of proteins across the endoplasmic reticulum. The T cells express the classical T cell genes TCR, CD3 and Interleukin-7 receptor (Il7r). Expression of the cytotoxic protease granzyme B (GZMB_1_) and CD8α and CD8β confirm this population as cytotoxic CD8^+^ T cells. CD8α and CD8β are lowly expressed by only a small percentage of cells suggesting that the transcription of these genes may occur in short bursts or there is a low cell surface expression level. A small group of lymphocytes were differentially clustered from the majority of T and B cells. These cells highly differentially expressed many histone genes, such as histone H2A, marker of proliferation Ki-67 (MKI67) and proliferating cell nuclear antigen (PCNA). This set of genes has greatest expression in actively proliferating cells, and we therefore named these cells proliferating lymphocytes. Some proliferating lymphocytes are more closely aligned to T cells (97 cells), while others are more closely aligned with the B cells (44 cells).

Two additional lymphocyte-related populations were found. Cluster 10, a rare population found in both the PBL and the spleen, is characterized by the expression of GATA-3 and several cytotoxic enzymes, including granzyme (GZM) B_2_, K and A. GATA-3 is a transcriptional activator that is shown to play a crucial role in the development of T cells, and specifically acts as a master regulator of T helper 2 (Th2) differentiation in mammalian species (Lee, Fields et al. 2001, Yagi, Zhu et al. 2011). We named this population cytotoxic GATA3+ cells.

The cluster labelled as NK-like cells express a somewhat unclear pattern of genes and it is hard to confidently assign an identify based on classical mammalian markers. These cells differentially express carcinoembryonic antigen-related cell adhesion molecule 6 (CEACAM6), killer cell lectin-like receptor subfamily B (KLRB1), nuclear factor erythroid 2 (NFE2), Colony Stimulating Factor 2 Receptor Alpha Subunit (CSF2RA) and the CD163 molecule. CEACAM6 is involved in cell adhesion, KLRB1 is a well-known NK receptor (Bennett, Zatsepina et al. 1996, Konjevic, Martinovic et al. 2009), NFE2 is a transcription factor, CSF2RA is a cytokine receptor, and CD163 is a hemoglobulin scavenger receptor. Collectively, these markers have been affiliated with a range of mammalian cell populations including neutrophils, T cells, NK cells, macrophages and granulocytes (Lanier, Chang et al. 1994, Skubitz and Skubitz 2008). Considering the combined expression profile of these genes, we putatively have named them NK-like cells.

The neutrophils represent the second largest cell population, and were identified by the expression of the cytotoxic genes eosinophil peroxidase (EPO) and non-specific cytotoxic cell receptor protein 1 (NCCRP1), genes involved in phagocytosis, neutrophil cytosolic factor 1 and 2 (NSF1, NSF2), and chemoattractant genes such as C-C motif chemokine 4 (CCL4). The gene classified as EPO has a sequence similar to both that of myeloperoxidase, the classic marker of neutrophils, and EPO. The most highly expressed gene of the neutrophils is low choriolytic enzyme (LCE), a proteolytic enzyme most commonly associated with egg hatching. Zebrafish neutrophils have been shown to express high choriolytic enzyme (HCE, also known as nephrosin) (Di, Lin et al. 2017), a related protein with high sequence similarity.

Erythrocytes were identified by the high expression of multiple haemoglobin genes (HBB). Erythrocytes also expressed immune related genes, such as NACHT, LRR and PYD domains-containing protein 12 (NLRP12), a potent mitigator of inflammation (Normand, Waldschmitt et al. 2018). Expression of thrombospondin-1 (THBS1), platelet glycoprotein Ib alpha chain (GP1BA) and thrombopoietin receptor (MPL) genes was used to identify the thrombocytes.

Macrophages were identified by the marker genes macrophage colony-stimulating factor 1 receptor (M-CSF1R), CSF2RA, and macrophage mannose receptor 1-like (MRC1). This cell population expresses many genes involved in the complement system, such as properdin (CFP) and complement factor B (CFB), and chemotactic genes, such as C-C motif chemokine 20 (CCL20).

Cluster 9 is a small population of cells found mostly in the spleen which express many innate immune genes including toll-like receptor 22 (TLR22), the chemokine receptor chemokine XC receptor 1 (CXCR1), bactericidal permeability-increasing proteins (BPI) and complement genes such as complement C1q-like protein 2 (C1QL2). We also observe the expression of the cytokine receptor fms like tyrosine kinase 3 (FLT3) and the transcription factor zinc finger 366 (ZNF366, also known as DC-SCRIPT), which have both been linked to differentiation of DCs in mammalian systems as well as in other teleost species (Zoccola, Delamare-Deboutteville et al. 2015). Another highly expressed gene in this cluster is allograft inflammatory factor 1 (AIF1), a gene which has recently been described in DCs (Elizondo, Andargie et al. 2017). Based on these markers, we have putatively named cells in this cluster DCs.

The cells tentatively named spleen stromal cells express many genes involved in fat metabolism and tissue structure, such as caveolae-associated protein 1 (CAVIN1), fatty acid-binding protein (FABP), apolipoprotein E (APOE). These cells are almost exclusively found in the spleen (94%). The expression profile of these cells is not as clearly differentiated as other clusters, as demonstrated in the heatmap (Figure 1B), with a variable expression of genes also found in other clusters. It is possible this cluster of cells is not a ‘true’ cell population and is merely an artefact of ambient RNA captured by beads in empty droplets. A small population of cells branching off from the spleen stroma, population 13, express markers for cell endothelium, including common lymphatic endothelial and vascular endothelial receptor-1 (CLEVER-1), a protein that is primarily expressed on high endothelial venules and lymphatic vessels (Irjala, Elima et al. 2003) where it supports the adhesion and transmigration of lymphocytes (Salmi, Koskinen et al. 2004), and plasmalemma vesicle associated protein (PLVAP), an endothelial cell-specific membrane protein (Stan 2008).

The cells from the unsorted spleen and the PBL samples are found in each cell cluster (Figure 2). The largest cell population of the unsorted spleen sample are the spleen stroma cells (27%), classified within the ‘other cells’ in the pie chart, followed by the cytotoxic T cells (22%), the B cells (16%), and then the erythrocytes (13%). In the PBL sample, B cells are the largest cell population (42%), followed by the T cells (25%) and neutrophils (18%). Cells from the sorted spleen S3 sample are populated mostly by neutrophils (35%), macrophages (28%) and erythrocytes (26%). The sorted blood B1 sample contains mostly thrombocytes (34%) and erythrocytes (23%), followed by cytotoxic T cells (19%) and B cells (19%).

Identification of cells by their transcriptional fingerprint is consistent with the characterisation of the major cell populations by microscopy (Figure 3). HE stained S3 sample, which was sorted as a relatively large and granulated population of spleen cells, revealed an apparent maturation of monocytes into macrophages, with a development from a lobed nucleus and few vacuoles to larger nuclei and more vacuoles. Vacuoles could also be seen in the erythrocytes, neutrophils and in some of the lymphocytes, supporting previous assertations that these are phagocytic cell types. Peroxidase staining of S3 sample revealed peroxidase positive myeloid cells, the neutrophils, and some peroxidase negative myeloid cells, the monocytes/macrophages. The ratio of peroxidase positive and peroxidase negative myeloid cells observed in the S3 population is similar to the ratio of neutrophils and macrophages seen in figure 2.

We next looked at the overall transcriptional activities of the different cell clusters (Figure 4A). The average number of transcripts per cell was noticeably high (1915 transcripts per cell) in the plasma cell group, whereas it was markedly low in the thrombocyte cluster (<500 transcripts per cell). DCs and macrophages also demonstrate a high transcriptional activity (1200-1300 transcripts per cell). Figure 4B shows the differential expression of cathepsin genes (CTSB and CTSL1) and an Fc-receptor gene (FCGR1a) in these two cell types. MHCI expression is present in all of the cell types as expected, however poor mapping has resulted in an inconclusive pattern across the cell clusters (supplementary figure 7).

## Discussion

This study, using state of the art sequencing technology on a non-model system, has provided a detailed molecular and cellular characterisation of the Atlantic cod immune system. The acquired knowledge will be highly beneficial for the development of antibodies towards cod-specific cell markers, our understanding of alternative vertebrate immune systems and potentially aid cod aquaculture and stock management.

Atlantic cod are able to mount a protective and specific immunity after vaccination (Lund, Bordal et al. 2007, Caipang, Brinchmann et al. 2009, Gudmundsdottir, Magnadottir et al. 2009, Magnadottir, Gudmundsdottir et al. 2009, Mikkelsen, Lund et al. 2011). Interestingly, this protective immunity is poorly correlated with specific antibody responses (Magnadottir, Jonsdottir et al. 2001, Mikkelsen, Lund et al. 2011). Indeed, based on these and other molecular observations, it was hypothesised that cod could lack functional MHCII molecules (Pilström, Warr et al. 2005) before it was definitely shown by the first complete cod genome. Despite major genetic losses within the CD4+ pathway and the limited response of specific antibody upon immunisation, in its natural environment cod are not particularly prone to infectious diseases (Pilström, Warr et al. 2005). How cod fights bacterial infections and how it acquires immunological memory, are puzzling questions that are both interesting scientifically and important practically for the cod aquaculture industries. With the near complete lack of antibody-based reagents and immortalised cod immune cell lines these mechanisms have not been fully understood, although some insights have been gained by genome data and transcriptome analyses at the whole organism or organ level (Solbakken, Jentoft et al. 2019a, Solbakken, Jentoft et al. 2019b). To find the exact immune cell composition and the steady-state gene expression of each cell subset is the first step toward a more detailed molecular mapping of the cod immune system.

This paper presents an overview of the immune cells found in the Atlantic cod peripheral blood and spleen, using single cell RNA sequencing and microscopy of both unfractionated and sorted cell populations based on size and granularity. By using unbiased clustering of global transcriptomics, we find thirteen different cell populations, with cell identity assigned to these populations based on their unique transcriptional profiles compared with known profiles from mammalian systems. Given the limited number of genes detected in each cell, we have restricted out interpretation of the data to assign cell population identities, without a more in-depth exploration of the cellular functions of these cell subsets. We identified the known major populations in the literature. These include: the cytotoxic T cells, B cells, erythrocytes, thrombocytes, neutrophils and macrophages. Overall, these six major cell populations make up 98% of hematopoietic cells in the spleen and PBL. In the lymphoid lineage, we identified sub-populations of both B and T cells that are actively dividing. In addition, we identified plasma cells as a separate subset in Atlantic cod, supporting old data where the presence of plasma cells was suggested by in situ hybridisation with immunoglobulin probes (Stenvik, Schroder et al. 2001). To note, a small subset of cells in this cluster were from PBL samples, therefore it is possible that this cluster also contains plasmablasts. Terminally differentiated plasma cells are rarely found in circulation, unlike the more immature plasmablasts. This data indicates that cod B cells have the capability of end-differentiation into plasma cells despite the lack of CD4+ T helper cells. Future studies are needed to clarify signalling pathways that are involved in B cell differentiation in cod.

Besides erythrocytes and thrombocytes, based on the top differentially expressed genes, we could clearly delineate the two major phagocytic and cytotoxic subsets within the myeloid lineage; namely macrophages and neutrophils. In mammalian systems, macrophages are important producers of cytokines as well as being antigen-presenting cells, especially in the spleen and lymph nodes. However, we could only find a handful of chemokines and no cytokine transcripts in our data. The low sensitivity of detecting transcript for single gene in any given single cell, combined with cytokines being expressed in short bursts only upon activation (Fang, Xie et al. 2013), could explain this particular pattern in our study.

The average number of transcripts that a cell clusters expresses may indicate, with a broad brushstroke, how active the cells are in steady state. Cells which are producing a lot of proteins, proliferating or carrying out multiple tasks, for example phagocytosis and antigen presentation, may be expected to have a higher transcript count than cell populations with fewer “tasks”. Unsurprisingly plasma cells are the most transcriptionally active cell population, with an average transcript count of 1915 detected transcripts, as this population of cells will be actively producing antibodies. The erythrocytes, neutrophils, NK-like cells, T cells, B cells and cytotoxic GATA3+ cells have medium levels of transcriptional activity, with approximately 650-1070 transcripts per cell. These levels represent most likely the steady-state transcriptional activity. The low level of transcriptional activity in thrombocytes is in accordance with the largely absent cytoplasm of these cells.

The proliferating lymphocytes, DCs and macrophages have a recorded transcript number of roughly 1200-1300 per cell. The proliferating lymphocytes are actively dividing so a high transcript number is anticipated. Teleost DC and macrophages are known to be phagocytic. A higher transcript number in these cells compared to the other known phagocytes - the neutrophils, erythrocytes and B cells, suggests that they may have additional tasks. The macrophages and the DCs are shown to express cathepsin genes (CTSB and CTSL1) and high affinity immunoglobulin gamma Fc receptor I A (FCGR1a). The gene annotation of FCGR1a is misleading since teleost does not have IgG. However, as annotation is based on sequence similarity to annotated genes in other organisms, it is probable that this gene represents an Ig-Fc binding receptor that could play a role in antibody-mediated uptake of antigens. Cathepsin B and L1 are lysosomal cysteine proteases that plays a major role in catabolism of proteins and thus a function in the processing and presentation of antigens via MHC (Hsieh, deRoos et al. 2002, Chapman 2006). The expression of genes involved in antigen presentation coupled with a high transcript number suggests the DCs and the macrophages may act as antigen presenting cells (APCs) in the Atlantic cod. It would be interesting to see if MHCI gene expression is higher in these cell populations, and if the expression increases following immune challenge. An initial analysis of MHCI expression in our data is inconclusive, mainly due to the low mapping efficiency. Dedicated efforts using longer reads and tailored bioinformatical tools that can deal with the complexities of MHCI, are needed.

Interestingly, the Atlantic cod has evolved MHCI expansion and an unusual repertoire of TLR receptors (Star, Nederbragt et al. 2011, Solbakken, Torresen et al. 2016). In addition, novel combination of endosomal sorting motifs was suggested to facilitate a more versatile use of MHCI through cross-presentation, and a potential MHCII-like functionality (Star, Nederbragt et al. 2011, Malmstrom, Jentoft et al. 2013). Whether the Atlantic cod is able to mount a cellular immune response that functionally resembles the T helper cells is unknown, and remains an intriguing issue for all the Gadiformes.

In our data, the expression of GATA-3 in a small subset of cells, with an expression profile that indicates close resemblance to the cytotoxic T cells is an interesting finding. GATA-3 has been identified and isolated from several species of teleost fish, including zebrafish (Neave, Rodaway et al. 1995), carp (Wang, Shang et al. 2013), salmonids (Kumari, Bogwald et al. 2009) and in Atlantic cod (Chi, Zhang et al. 2012). GATA-3 expression was detected in surface-IgM-negative lymphocytes in carp (Takizawa, Mizunaga et al. 2008). Interestingly, the expression of GATA-3 in Atlantic cod was shown to be increased following stimulation by the T cell-stimulant phorbol 12-myristate 13-acetate (PMA) (Chi, Zhang et al. 2012), suggesting the presence of GATA-3 in activated T cells. In mammals, the transcription factor GATA-3 plays an essential role in CD4+ T cell development and survival, and is necessary for the differentiation of naive CD4+ T cells to T helper (Th) 2 cells (Murphy and Reiner 2002, Bosselut 2004, Ho and Pai 2007). However, classical T helper cells are absent in Gadiformes and this cytotoxic GATA3^+^ cluster lacks the cytotoxic T cell markers seen in our data, suggesting that this population may belong to a different lineage than T cells.

In mammals GATA-3 is also central to the development of innate lymphoid cells (ILCs), chiefly the ILC2 lineage (Hoyler, Klose et al. 2012, Tindemans, Serafini et al. 2014). Recently, it was shown that GATA-3 expression was also important for ILC2 in zebrafish (Hernandez, Strzelecka et al. 2018). ILC2 cells are also known as innate helper 2 cells (Moro, Yamada et al. 2010) based on similar cytokine secretion profile. Thus, in summary, the cells described here possibly represent a form of helper ILC. At the same time, these cells also show granzyme expression; indicating possible dual cytotoxic and helper functions. Future studies should look into how this small but intriguing cell subset behaves during immune perturbation, such as immunisation and infection.

In conclusion, our results provide the most complete overall mapping of the repertoire of Atlantic cod immune cells present to date. In addition to describing in more detail the major cell subsets, we also describe for the first time in Atlantic cod cells that may represent DCs, NK-like cells and ILCs, as well as suggest that macrophages and DCs may act as APCs. This work provides an expression profile baseline of the Atlantic cod in a steady state, which provides a foundation for future work with immune system challenge experiments.

## Acknowledgements

Ketil Hylland- providing cod

Marine Brieuc- bioinformatics support

Ole Tørresen - bioinformatics support

Norwegian Sequencing Centre (NSC)

Espen Bækkevold – help with histology

UiO – Convergence Environment COMPARE

