## Supplementary for "Single cell transcriptome profiling of the Atlantic cod immune system"

Supplementary figures

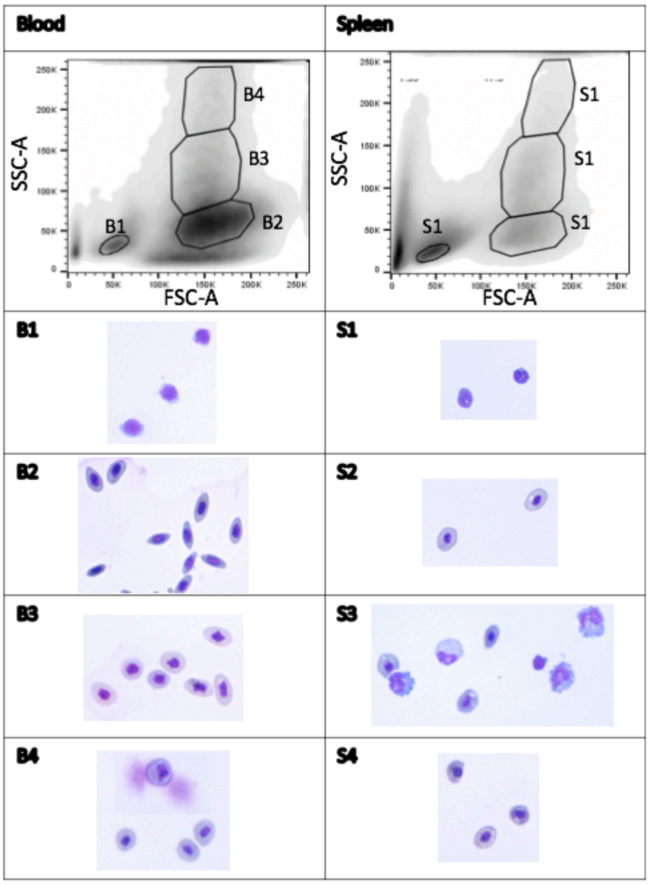


Supplementary figure 1. Fresh Atlantic cod blood and spleen samples were flow-sorted into four populations each based on forward scatter and side scatter gating. Representative images of HE-stained cells from blood sub-populations (B1-B4) are shown on the left, while spleen sub-populations (S1-S4) are shown on the right. The percentage of the populations are as follows: B1-3%, B2-28%, B3-13%, B4-4%, S1-18%, S2-8%, S3-8%, S4-1.5%

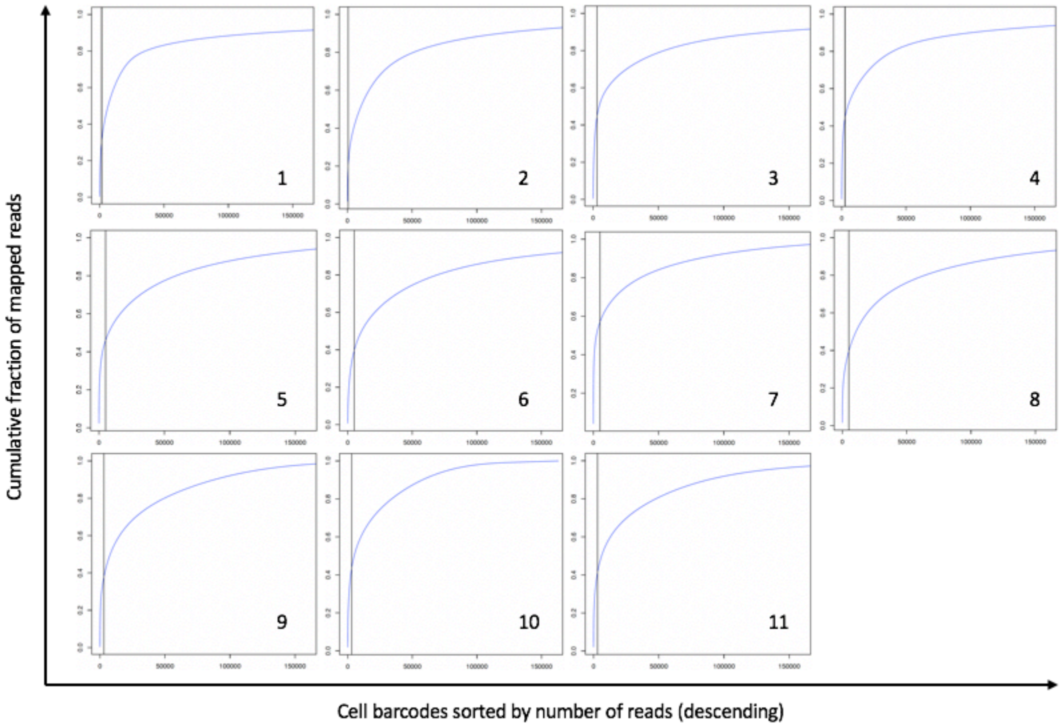


Supplementary figure 2. Cumulative distribution of reads used to identify STAMPs in a pool of amplified beads. Each number, 1-11, represents the sample identity. Drop-seq involves the generation of single-cell RNA profiles by adding a single cell to a single droplet containing a single bead. The large majority of amplified beads are not encapsulated with a cell and so are not exposed to a single cell's RNA, only the ambient RNA present in solution following droplet breakage. To identify the cell barcodes which do correspond to STAMPs, cell barcodes from each sample used in the experiment are arranged in decreasing number of reads, and the cumulative fraction of reads is plotted. The predicted number of individual cells calculated from the counted and aliquoted beads for each sample was estimated during the laboratory procedure. However, there is no clear inflection point at these estimated numbers. To be sure to include all salient information it was decided to include the first 600-3000 cell barcodes (shown by the black line), to be further filtered at later steps.

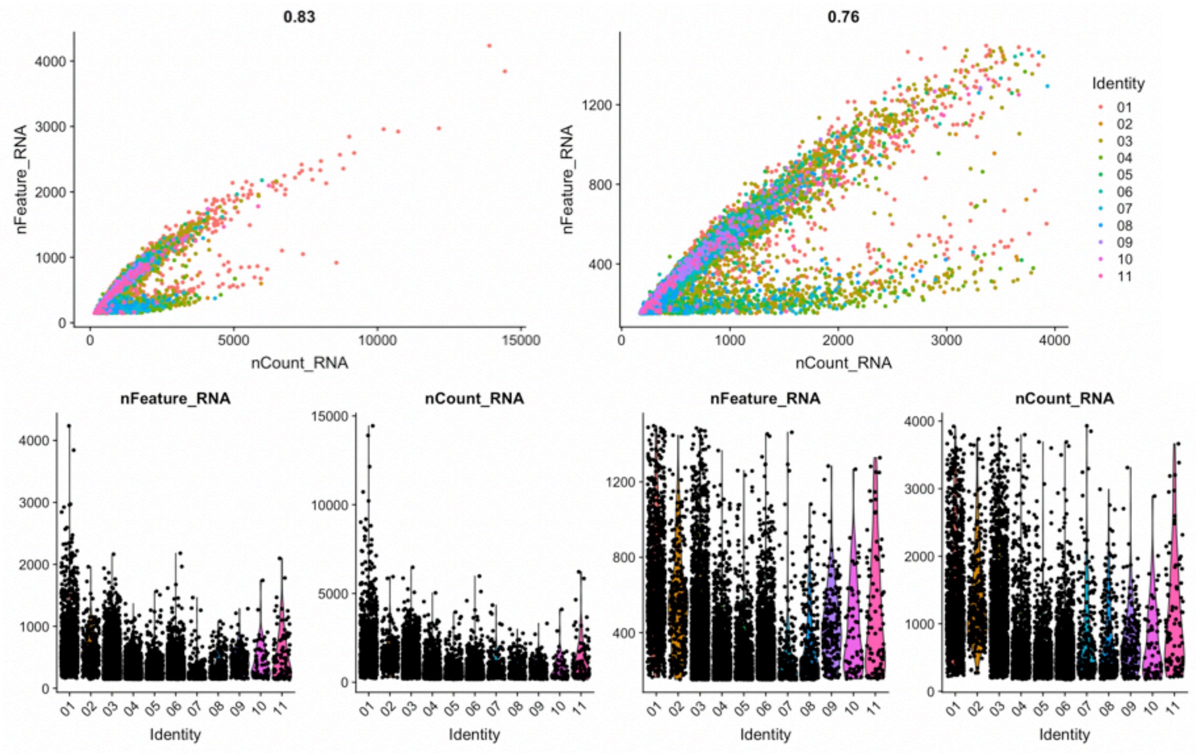


Supplementary figure 3. Feature scatter (top) and violin plots (bottom) showing the 11 samples before (left) and after (right) quality control cut offs are applied. Cells with a gene count (nFeature_RNA) of fewer than 150 and more than 1500, cells with a total number of molecules of more than 4000 (nCount_RNA) were filtered away in order to remove low-quality cells and possible cell multiplets.

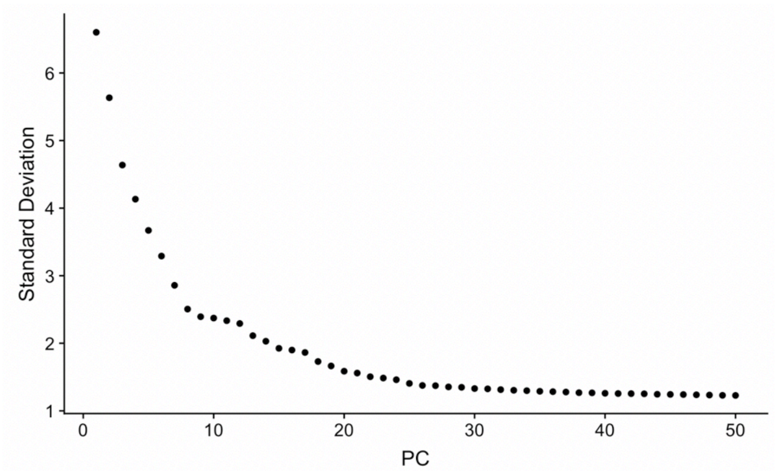


Supplementary figure 4.
Elbow plot showing a ranking of principle components based upon the percentage of variance explained by each. Here we observe an 'elbow' around PC25-30, suggesting that the majority of true signal is captured in the first 30 PCs.

| | Sample number | Tissue | Library size (millions of fragments) | Average mapping (%) | Reads mapped (millions) | Number of cells included after filtering | Average number of mapped reads per cell | Average number of mapped transcripts per cell* | Average number of mapped genes per cell |
|---|---|---|---|---|---|---|---|---|---|
| **Fish 1** | 1 | Spleen | 114 | 53 | 61 | 1153 | 4157 | 1356 | 628 |
| | 2 | Spleen | 51 | 57 | 29 | 327 | 5922 | 1381 | 557 |
| | 3 | S3 sorted spleen | 116 | 55 | 63 | 2102 | 6083 | 1059 | 430 |
| | 4 | B1 sorted blood | 60 | 54 | 33 | 1226 | 4985 | 656 | 318 |
| **Fish 2** | 5 | Spleen | 66 | 59 | 39 | 758 | 5333 | 637 | 306 |
| | 6 | PBL | 60 | 59 | 35 | 1780 | 2120 | 583 | 340 |
| | 7 | S3 sorted spleen | 15 | 59 | 9 | 230 | 2610 | 793 | 283 |
| | 8 | B1 sorted blood | 23 | 59 | 14 | 256 | 2794 | 730 | 332 |
| | 9 | P1 sorted PBL | 18 | 59 | 10 | 197 | 6882 | 753 | 437 |
| | 10 | P2 sorted PBL | 7 | 59 | 4 | 60 | 5021 | 731 | 418 |
| | 11 | P3 sorted PBL | 22 | 56 | 12 | 91 | 5231 | 1113 | 503 |
| | | | Total = 552 | Average = 57 | Total = 309 | Total = 8180 | **Weighted average = 4695 | **Weighted average = 884 | **Weighted average = 409 |

**Supplementary table 1.** Overview of sample origin and sequencing results. Blood and spleen sample were taken from two Atlantic cod. An average of 57% of sequencing reads were mapped to the Atlantic cod genome (gadmor3). Following drop-seq computational protocol pipeline and the Seurat package filtering steps gene data from 8180 cells are included, with an average of 409 genes expressed per cell. *Number of transcripts per cell given are after accounting for PCR amplification **Averages are weighted by number of cells per sample.

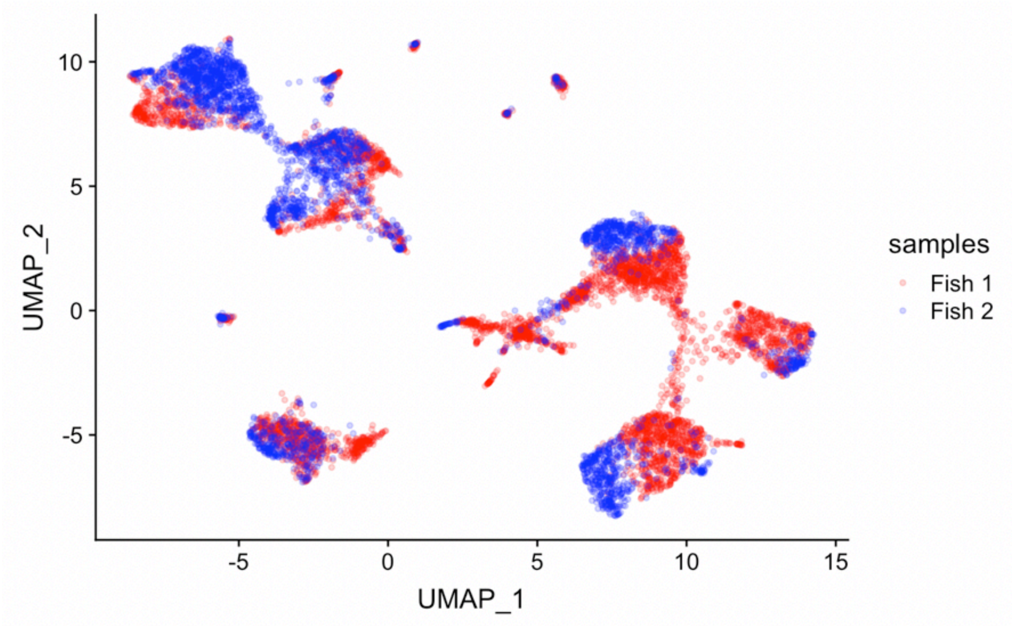


Supplementary figure 5.
Plot showing the distribution of cells from Fish 1 (sample 1-4) and fish 2 (samples 5-11) across the clusters.

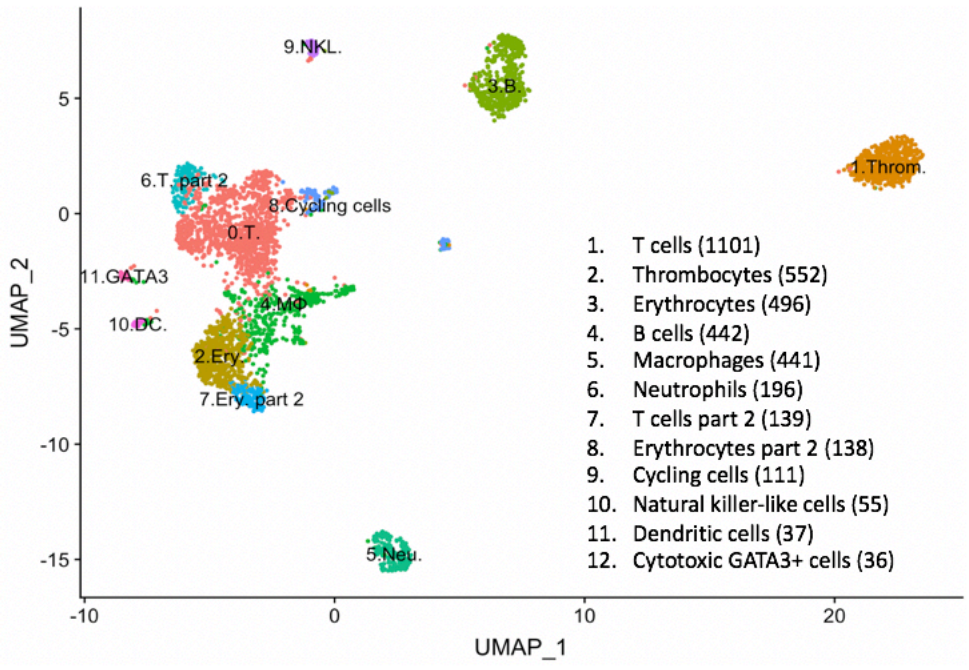


Supplementary figure 6. Cell types from unsorted blood and spleen visualised in two dimensions using UMAP. Cells are from pilot study using wild-caught Atlantic cod in Oslo Fjørd. Putative cell cluster labels are based upon the differential gene expression of known markers in mammals. The same parameters used in the main study have been applied: cells with a gene count of fewer than 150 and a gene count of more than 1500, and cells with a total number of molecules of more than 4000 were filtered away in order to remove low-quality cells and possible cell multiplets. Genes expressed in less than 5 cells were excluded. 30 principal components were included and the resolution for clustering was 0.35. The T cells and erythrocyte cell clusters are split into two subclusters at this resolution due to batch effects.

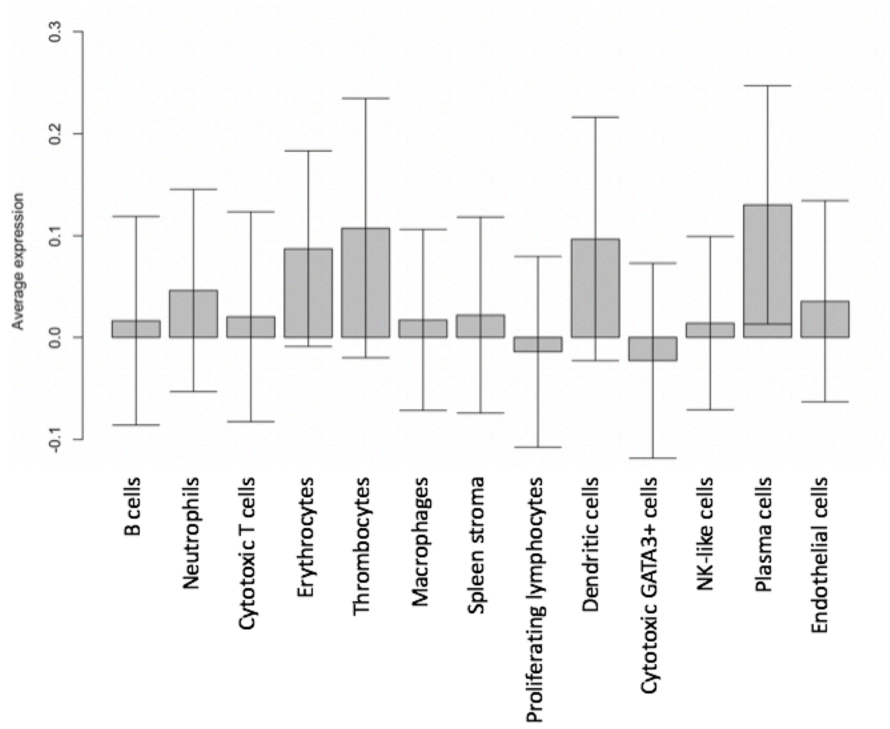


Supplementary figure 7. Average expression of MHCI genes across each cell cluster. Error bars show standard deviation.
